# Characterization of α_2_-Adrenergic Receptors Mediated Vasoconstriction in *ex vivo* model

**DOI:** 10.1101/2021.04.23.441223

**Authors:** Molly Yao

## Abstract

**Introduction:** rat tail serves as a thermoregulatory organ by dilating or constricting tail blood vessels and rat tail lateral veins play a role in cutaneous circulation. However, this cutaneous vessel has never been examined and evaluated as a potential candidate *ex vivo* model to study peripheral vascular diseases. This study aims to investigate vascular tone mediated through α_2_-adenergic receptor on rat tail vein under different temperatures.

**Methods:** Vasoconstriction stimulated by α_2_-adrenergic receptor selective agonists UK14304, guanabenz was thoroughly examined in isolated rat tail lateral vein. Susceptibility of cutaneous vessel to temperature changes was investigated under 37 °C and 28 °C. Differentiated vascular reactivity exhibited at different temperature was further explored with different α_2_-adrenergic receptor selective agonists and validated with antagonism by α_1_-adrenergic receptor and α_2_-adrenergic receptor selective antagonist prazosin and RX821002, respectively.

**Results:** Vascular tone of freshly isolated vein remained same compared with that of stored in pre-gassed physiologic buffer at 4 °C overnight. Presence of endothelium and repeated administration of UK14304 did not alter contractile property. Vessels away from torso (distal) showed significantly different contractile character compared with portion close to torso (proximal) at 28 °C but kept uniform at 37 °C. Enhanced vasoconstriction along with increased potency of α_2_-adrenergic receptor agonists UK14304 and guanabenz was consistently present at 28 °C, independent of temperature change orders. Potentiated vasoconstriction present at 28 °C was later proved via α_2_-adrenergic receptor alone.

**Discussion:** Harvest and preparation of rat tail lateral vein was described in details. Contractile property of venous preparations stimulated by α_2_-adrenergic receptor selective agonists was first time examined. Differentiated vasoconstriction at moderate cooling temperature was described and confirmed to through α_2_-adrenergic receptor activation. Rat tail lateral vein is found valuable for studies of cutaneous circulation altered by temperature changes.

## Introduction

The rat tail is an essential thermoregulatory organ based on its specific features, e.g. lack of fur, a large ratio of surface area to volume, and a unique vascular organization. The rat tail regulates body temperature by dilating or constricting tail blood vessels. *In-vivo* study of blood vessels in the rat tail revealed a significant diameter enlargement of the tail veins compared to the tail arteries in response to increase in body temperature (G. Vanhoutte, Verhoye, Raman, Roberts, & Van der Linden, 2002). Inverse, excessive heat loss is essentially prevented by decreasing tail cutaneous blood flow in response to local or whole body cooling, which is mediated by increased sympathetic tone as well as locally regulated vasoconstriction (Flavahan, 1991; G. Vanhoutte et al., 2002). Similarly, arteriovenous anastomoses in human digits are under control of the central nervous system and the local temperature variations. Arteriovenous anastomoses regulate the skin temperature through volume adjustments in the superficial venous bed (Vangaard, 1948). These similarities in anatomy and function establish the rat tail as a valuable *in vitro* model to better understand characteristics of cutaneous blood vessels in thermoregulation. Raynaud’s phenomenon is characterized by exaggerated cutaneous vasospasm in response to environmental cold temperature or emotional stress (Wigley, 2002). Recurrent cycles of excessive vasoconstriction-induced transient ischemia followed by reperfusion results in reactive oxygen species generation leading to ischemia-reperfusion-injury (IRI) (Cooke & Marshall, 2005; Liu et al., 2013; Simonini, Pignone, Generini, Falcini, & Cerinic, 2000). Severity of IRI varies from superficial ulceration to necrosis of deep tissue with gangrene and amputation may necessitate. Similar to Raynaud’s phenomenon occurred to human, ringtail, also known as tail necrosis, is an epidermal disease as a consequence of annular constrictions along its length that may occur in rodents like rats and mice. In the most severely affected animals, the tail may become gangrenous and drop off.

A variety of studies on the thermoregulatory function of cutaneous or subcutaneous blood vessels have been performed in the canine saphenous vein (Flavahan, Lindblad, Verbeuren, Shepherd, & Vanhoutte, 1985; Flavahan & Vanhoutte, 1986; Janssens & Vanhoutte, 1978), equine digital vein (Zerpa, Berhane, Elliott, & Bailey, 2007), rabbit ear and femoral artery (Garcia-Villalon et al., 1992), mouse tail artery (Chotani, Flavahan, Mitra, Daunt, & Flavahan, 2000) and rat tail artery (Jantschak et al., 2010; Souza, Padilha, Stefanon, & Vassallo, 2008). Histological examination of the rat tail found one mid-ventral artery enclosed in a groove covered by fascia and two cutaneous lateral veins (Staszyk, Bohnet, Gasse, & Hackbarth, 2003). Anatomy of the rat tail vein suggests it might be highly susceptible to temperature fluctuation. However, vascular reactivity of rat tail lateral vein has not yet been characterized.

Norepinephrine released from sympathetic nerve ending activates postjunctional α_1_- and α_2_-adrenergic receptors (AR) in cutaneous vessels to cause vasoconstriction and reduce heat dissipation. The objective of this paper is to describe in detail a technique for reliably harvesting and quantifying vasoconstriction of rat tail lateral vein via α_2_-AR-mediated signaling transduction. It is anticipated that this approach should aid investigation of novel vessels from other tissue and species where similar considerations and problems apply.

## 2. Methods

### 2.1 Animals

All experiments were conducted under the regulations of the Institutional Animal Care and Use Committee of Creighton University and in compliance with USDA regulations. Male Sprague Dawley rats (250-300 g) obtained from Charles River Laboratories (Wilmington, MA) were housed in plastic cages in temperature-controlled room (20±2 °C, 50% humidity) on a 12-hour light-dark cycle (light on 8:00 to 20:00). They were allowed *ad libitum* access to a standard laboratory diet and water.

### 2.2 Chemicals and reagents

The drugs used were obtained from the following sources: L-phenylephrine HCl, acetylcholine Cl (Sigma-Aldrich, St. Louis, MO), UK14304 tartrate (Pfizer, Groton, Connecticut), guanabenz acetate, RX821002 hydrochloride (Sigma-RBI, Natick, MA). All drugs were dissolved in 0.9% saline then diluted to the desired concentration with KHS.

### 2.3 Solutions

Krebs-Henseleit solution (KHS) consists of 118.4 mM NaCl, 11.1 mM glucose, 25 mM NaHCO_3_, 4.7 mM KCl, 1.2 mM MgSO_4_, 1.2 mM KH_2_PO_4_, 2.5 mM CaCl_2_, and 0.029 mM Na_2_Ca EDTA.

### 2.4 Isolated rat lateral tail vein preparation

The animals were euthanized by exposure to carbon dioxide. The rat tail was fixed on a dissection platform with adhesive tape, marked one-third along the length of the tail from the base of the tail as proximal and one-third further towards the tail tip as middle. Rat lateral tail vein was exposed right after peeling off skin with a #10 scalpel. The abundant elastic tissues in the tunica media of arteries or arterioles render the opening of lumen constantly open post-dissection with blood draining. In contrast, it becomes technically impossible to relocate the opening of lumen in isolated rat tail vein after the same procedure, making mounting isolated vessel to organ bath impracticable. A 0.15 mm-diameter nylon monofilament was inserted into the lumen of the vein to keep it accessible for mounting. The veins were removed, transferred to a dissecting plated and immersed in KHS then cleared from surrounding connective tissue. Tail veins were cut into 3 mm-long ring segments. Ring segments were horizontally mounted between two L-shaped stainless steel pins passed through the vessel lumen. One pin was attached to a Grass FT.03 isometric tension transducer (Grass Instruments, Quincy, MA) for measurement of isometric tension while the other pin was held in a fixed position. Rings were placed in water-jacketed organ bath chambers filled with 10 ml KHS continuously gassed with 95% O_2_-5% CO_2_, pH 7.4, and maintained at 37 °C or 28 °C through heating circulating pump.

### 2.5 Contractile property of different portions of rat tail vein

The length of the rat tail vein ranged from 160 to 200 mm. Contractile property of proximal and distal portion from rat tail vein was measured. Specifically, the proximal portion was defined as starting from the bottom of the torso to one sixth the length of the tail. The distal portion was defined as beginning from the end of the first third to one half of the tail length from the torso.

### 2.6 Storage of isolated rat tail vein at 4 °C

Two lateral veins can be collected from each animal. Isolated rat tail veins clear of fat and connective tissue were stored in a flask containing Krebs solution pre-gassed with 95% O_2_-5% CO_2_. The flask was tightly sealed with parafilm and kept in the refrigerator for approximately 22 hrs. Contractile property of rat tail vein stored at 4 °C overnight was compared to that of freshly prepared.

### 2.7 Determination of optimal resting-tension

Distension of vessels with circumferentially arranged smooth muscles brings the actin and myosin fibers into better alignment, allowing optimal force to develop in response to a pharmacologic agent (O’Rourke, Blaxall, Iversen, & Bylund, 1994). A length-tension relationship was first examined to establish the passive tension at which rat tail lateral venous rings performed optimally under active stimulus. 60 mM KCl solution was chosen for this study because KCl induces contraction due to membrane depolarization leading to so called electromechanical coupling (Clarke & Harris, 2002). Fresh venous ring preparations were equilibrated in KHS at 37 °C for 1 hour without tissue stretching. During equilibration KHS was replaced every 20 minutes. Venous rings were challenged with 60 mM KCl, active force generation was recorded and then tissues were washed for 30 min. The passive tension placed on the tissues was increased to next level. This procedure was repeated with passive tension ranging from 100 to 400 mg. A passive-active tension curve was generated to determine the optimal resting tension.

### 2.8 Isometric tension measurement of UK14304-induced vasoconstriction

Rat tail venous ring segments were equilibrated in KHS for 1 hour at a resting tension of 300 mg determined in method detailed above. Ring segments were contracted with 60 mM KCl followed by washout and a 30 minutes relaxation. This step was repeated once. To assess the presence of endothelium, ring segments were contracted with 30-100 nM phenylephrine reaching a plateau followed by 1 μM of endothelium-dependent vasodilator acetylcholine. Ring segments were then thoroughly washed for 45 minutes with KHS.

#### 2.8.1 Repeated administration UK14304

As an internal control each venous ring was twice stimulated by the same agonist, i.e., in the absence and presence of antagonist. To determine spontaneous change due to repeated treatment by the same agonist at different time, multiple comparisons between repeated treatments of agonist to the same venous ring were performed. A cumulative concentration-response curve for UK14304-induced contraction was first generated, followed by drug washout and equilibration in drug-free KHS for 1 hr. Cumulative concentration-response curves for UK14304 were then repeated on the same tissue.

#### 2.8.2 Incubation temperature shift between 28 °c and 37 °C

Cumulative concentration-response curves for α_2_-AR selective agonist UK14304- or guanabenz-induced vasoconstriction were first constructed at 28 °C by increasing the agonist concentration cumulatively by half-log increments until a maximal response was obtained. The bath temperature was warmed up to 37 °C and venous rings were equilibrated for 1 hr. Cumulative concentration-response curves for UK14304 or guanabenz were generated and the rings were then washed with KHS till the tension returned to the resting tone. Reversed sequence of temperature shift was also examined. Specifically, temperature of the organ bath was cooled down from 37 °C to 28 °C and maintained at that temperature for 1 hr and UK14304- or guanabenz-stimulated concentration-response curves at both temperatures were repeated.

#### 2.8.3 UK14304 in the absence and presence of antagonist RX821002

α_2_-AR antagonist RX821002 was used to confirm UK14304-stimulated vasoconstriction at both 37 °C and 28 °C is mediated through α_2_-AR in the rat tail vein. Following equilibration and KCl contraction at either 37 °C or 28 °C, cumulative UK14304 concentration-response curves (1×10^−9^ M to 3×10^−5^ M) were obtained. Rings were washed with KHS, incubated with 50 nM of RX821002 for 1 hr, and cumulative UK14304 concentration-response curves were then repeated.

### 2.9 Effect of cooling on 60 mM KCl-stimulated contraction of rat tail vein

All of UK14304-induced vasoconstriction recorded in tension (mg) was normalized to percentage of vasoconstriction in response to 60 mK KCl. To determine effect of cooling from 37 °C to 28 °C on vasoconstriction induced by 60 mM KCl, rat tail veins were contracted with 60 mM KCl followed by washout and equilibration for 30 min. This procedure was repeated four times at 28 °C. Rings were then incubated for 1 hrs after the temperature of the organ bath was increased to 37 °C. The same procedure of KCl administration and washout described above was then repeated four times at 37 °C. This protocol was performed in duplicate or triplicate on tissues from a single animal. Contraction induced by the first administration of KCl was excluded from data analysis.

### 2.10 Calculation of equilibrium dissociation constants and data analysis

Agonist concentration-response curves (CRCs) were plotted, the maximal contraction and EC_50_ values (the concentration of agonist generating 50% of the maximal response) were calculated using all points on the CRCs using least sum of square nonlinear regression curve fitting with GraphPad Prism 5.0 (GraphPad Software, San Diego, CA).

When an antagonist causes parallel rightward shifts of CRC without change in the maximal responses, this antagonist is considered competitive. For each concentration-response curve for UK14304 or guanabenz, 2-way ANOVA followed by a Bonferroni’s multiple comparison test was performed, significant difference accepted at *p*<0.05.

The logarithm of the EC_50_ value was used in all statistical comparisons while these values are presented as arithmetic mean ± S.E.M for *n* animals. The unpaired student’s *t*-test was used for statistical comparisons with a *p*<0.05 level of probability accepted as a significant difference.

## 3. Results

### 3.1 Determination of optimal resting tension

A length-tension relationship was first performed to establish the passive tension at which rat tail lateral venous rings performed optimally under active stimulus. Fig. 1 depicts the findings that a passive tension of more than 300 mg results in an optimum and maximal contraction to KCl. A resting tension of 300 mg was used in all following experiments.

**Fig. 1.**
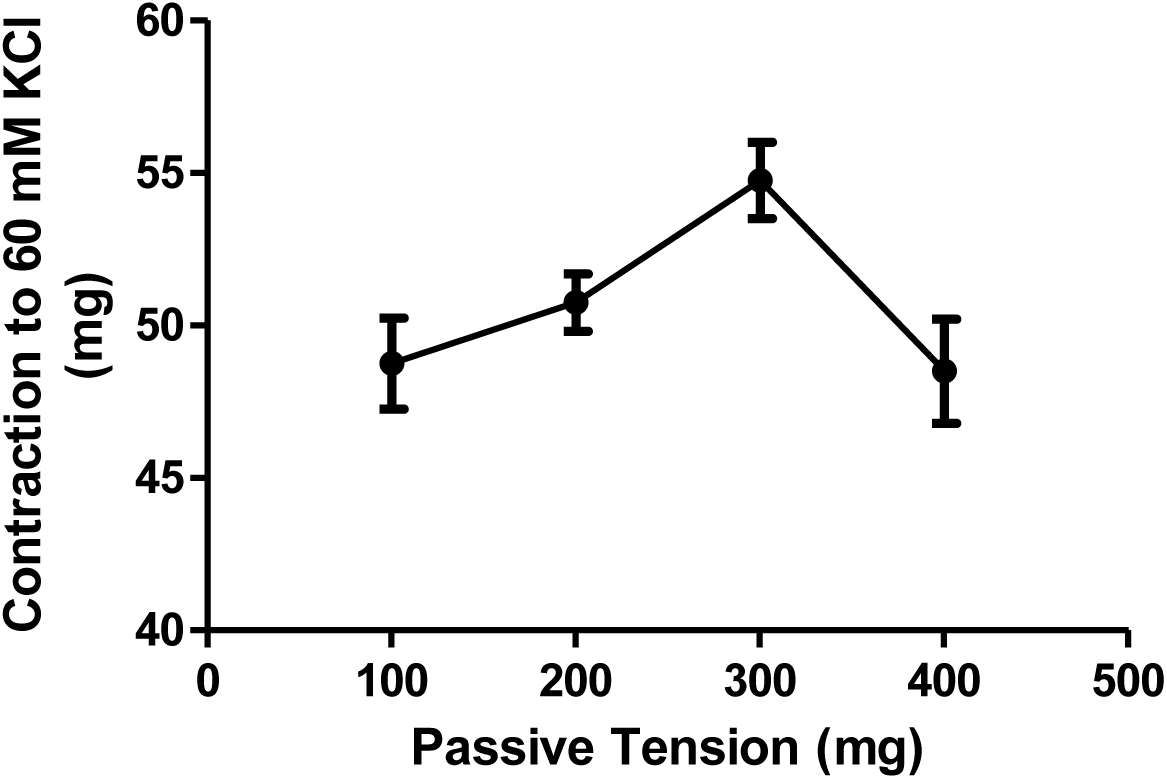
Active-length relationship for the isolated rat tail veins. Points represent means and vertical bars the S.E.M. for 4 experiments using individual rat tail veins, each taken from different animals.

### 3.2 Responsiveness of rat tail vein to α_2_-AR selective agonist UK14303

As showed in Fig. 2 through 6, isolated rat tail later vein generated robust contractions in response to UK14304 administration without requisite pre-contraction by either norepinephrine or serotonin as seen in rat tail artery (Jantschak et al., 2010), mouse tail artery (Chotani et al., 2000) or porcine pulmonary artery (Jantschak & Pertz, 2012).

**Fig. 2.**
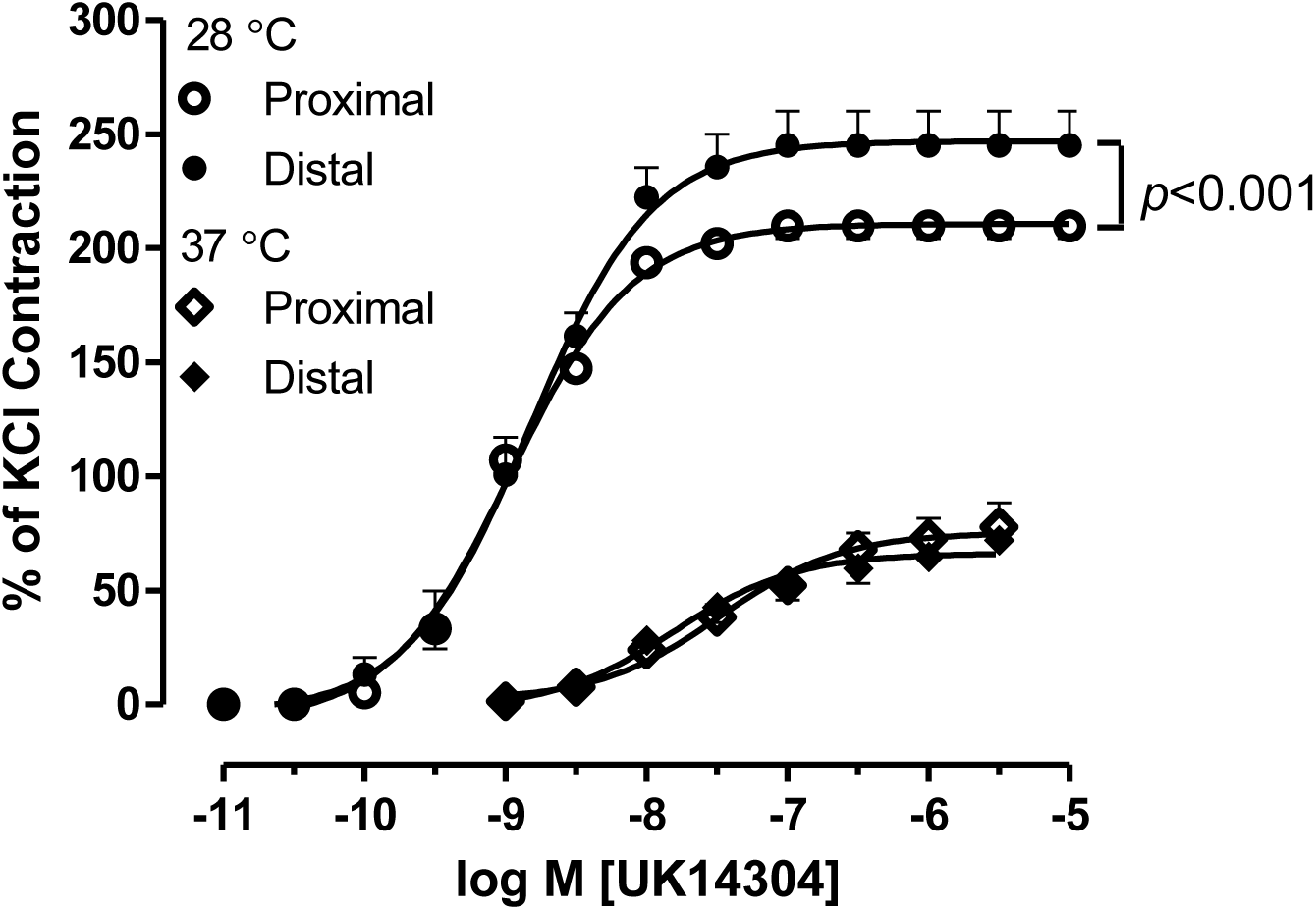
Effect of the distance of the rat tail vein ring from the torso on concentration-response curves for UK14304. Contraction generated by proximal and distal sections of rat tail veins at 37 °C and 28 °C are shown. Points are the mean ± S.E.M. of 5 experiments using individual rat tail veins, each taken from different animals.

### 3.3 Effect of the distance from the torso of the rat tail vein on contraction

Rat tail artery presented differentiated vascular tone at segments of base, middle and tail end (Morrow & Creese, 1986). Whether vasoconstriction of proximal portion is different from that of distal portion from the same tail vein was examine. It can be clearly seen in Fig. 2 that the distance from the torso did not affect UK14304-mediated contraction of the rat tail vein at 37 °C. As temperature is cooling down to 28 °C, CRC mediated by the proximal portion of rat tail vein is significantly different from the distal portion (*p*<0.001) without change in either potency or maximal contraction. Table 1 lists potencies and maximal contractions from these experiments.

**Table 1.** Summary of UK14304 potencies and maximal contractions. Potencies are listed as EC50 values and maximal contraction as percentage of 60 mM KCl-induced contraction. Each value is the mean ± S.E.M. of measurements made in number of animals present in the table.

|  | 37 °C |  |  | 28 °C |  |  |
| --- | --- | --- | --- | --- | --- | --- |
|  | EC<sub>50</sub><br>(nM) | Maximal Contraction<br>(% of KCl Contraction) | n | EC<sub>50</sub><br>(nM) | Maximal Contraction<br>(% of KCl Contraction) | n |
| Fresh | 58 ±21 | 77 ±9 | 4 | 1.3 ±0.3 | 190 ±13 | 6 |
| Stored | 51 ±17 | 85 ±10 |  | 1.6 ±0.4 | 210 ±11 |  |
| Proximal Sections | 46 ±21 | 78 ±11 | 5 | 1.1 ±0.1 | 210 ±5 | 5 |
| Distal Sections | 69 ±58 | 72 ±5 |  | 1.5 ±0.3 | 250 ±15 |  |
| + Endothelium | 7.3 ±1.0 | 81 ±8 | 4 | 1.2 ±0.2 | 190 ±19 | 6 |
| - Endothelium | 11 ±2 | 75 ±12 |  | 1.1 ±0.3 | 180 ±13 |  |
| 1<sup>st</sup> Treatment | 9.2 ±2.4 | 74 ±17 | 4 | 8.3 ±4.3 | 230 ±19 | 4 |
| 2<sup>nd</sup> Treatment | 9.2 ±4.2 | 63 ±14 |  | 10 ±6 | 240 ±19 |  |

### 3.4 Effect of endothelium on contraction of rat tail vein

α_2_-AR present on the endothelium may alter UK14304-mediated contraction of rat tail vein. UK14304 CRCs were compared for venous rings with either intact or denuded endothelium. Presence of the endothelium has no effect on UK14304-induced contraction (Fig. 3). Potencies and maximal contractions of veins with or without endothelium are summarized in Table 1.

**Fig. 3.**
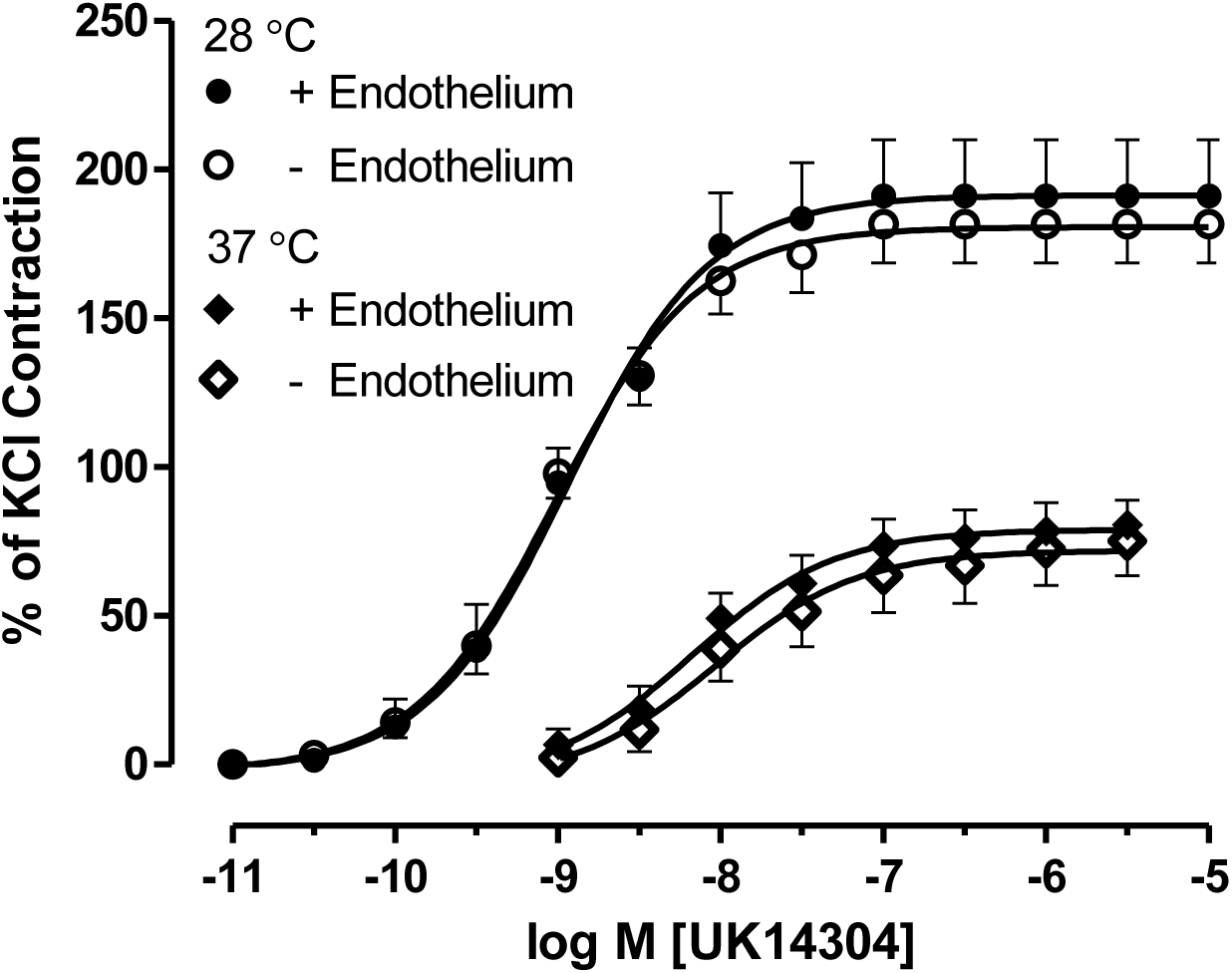
Effect of endothelium on concentration-response curves for UK14304 in rat tail veins. Veins with endothelium or without endothelium were studied at 37 °C and 28 °C. Points are the mean ± S.E.M. of 4-6 experiments using individual rat tail veins, each taken from different animals.

### 3.5 Effect of storage at 4 °C for 22 hrs on contraction of rat tail vein

Overnight storage of isolated tissue in physiological solution pre-gassed with combined O_2_-CO_2_ at low temperature, e.g. 4 °C, is a common and efficient way to maximize utilization of tissue samples also minimize animal number to be used. Contractile response to UK14304 in freshly prepared venous rings was compared to that in stored tissues. **Fig. 4** showed storage at 4 °C for overnight does not alter contractile response of rat tail veins, therefore isolated rat tail veins can be saved for next day use without altering contractile property to UK14304. Potencies and maximal contractions from these studies were listed in Table 1.

**Fig. 4.**
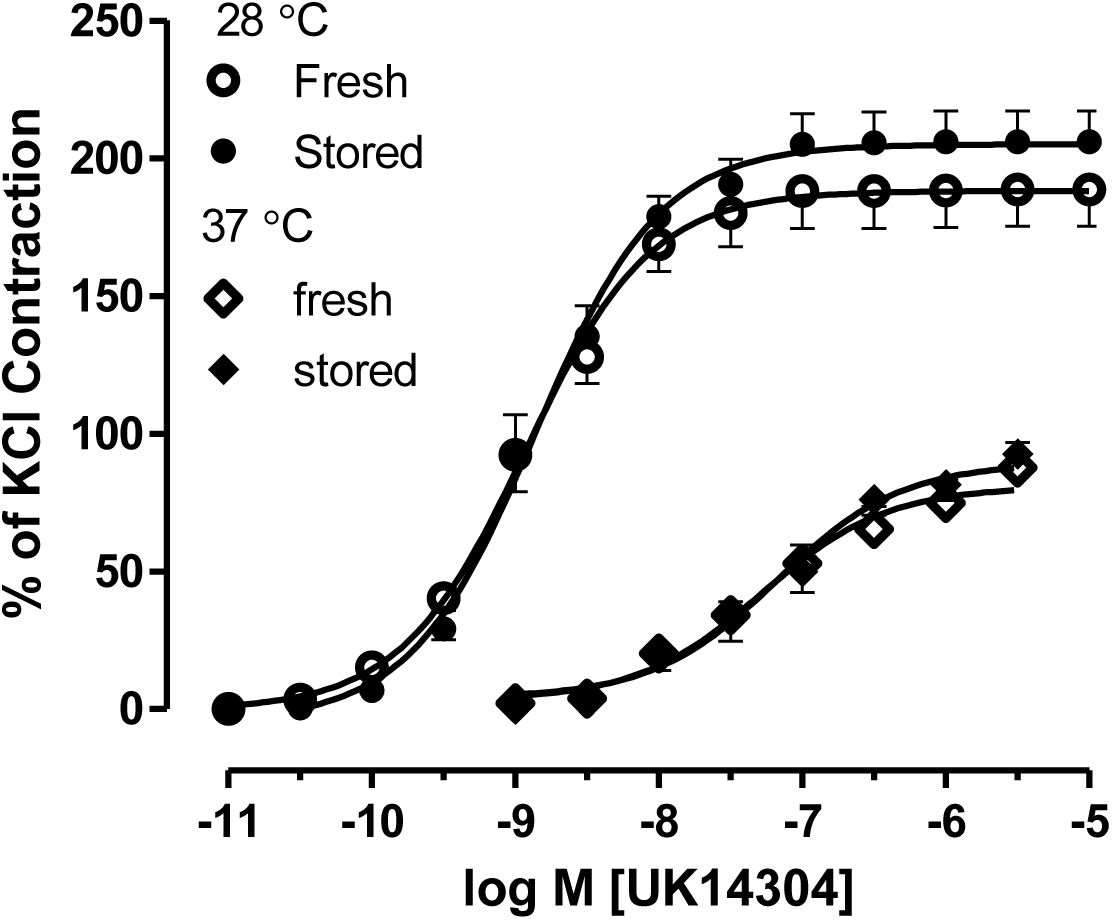
Effect of storage at 4 °C for 22 hrs on concentration-response curves for UK14304. Contraction generated by freshly prepared and stored rat tail veins at 37 °C and 28 °C are shown. Points are the mean ± S.E.M. of 4-6 experiments using individual rat tail veins, each taken from different animals.

### 3.6 Effect of repeated stimulation of rat tail vein by UK14304

Desensitization and consequent down-regulation induced by sustained or repeated agonist stimulation is a mechanism to prevent excessive activation shown in many receptor-effector systems, including α_2_-adrenoceptors (Flavahan, 2008; Saunders & Limbird, 1999). A venous ring can be considered as an internal control providing no significant change in contractile property when stimulated by UK14304 for multiple times. If desensitization occurred to the rat tail vein, internal control is not appropriate and a separate venous ring from the same animal must be used accordingly. Fig. 5 presented almost identical CRCs of the rat tail veins treated with UK14304 repeatedly with 1 hour apart. This study justifies validity of venous ring as internal control. Table 1 lists potencies and maximal contractions from these experiments.

**Fig. 5.**
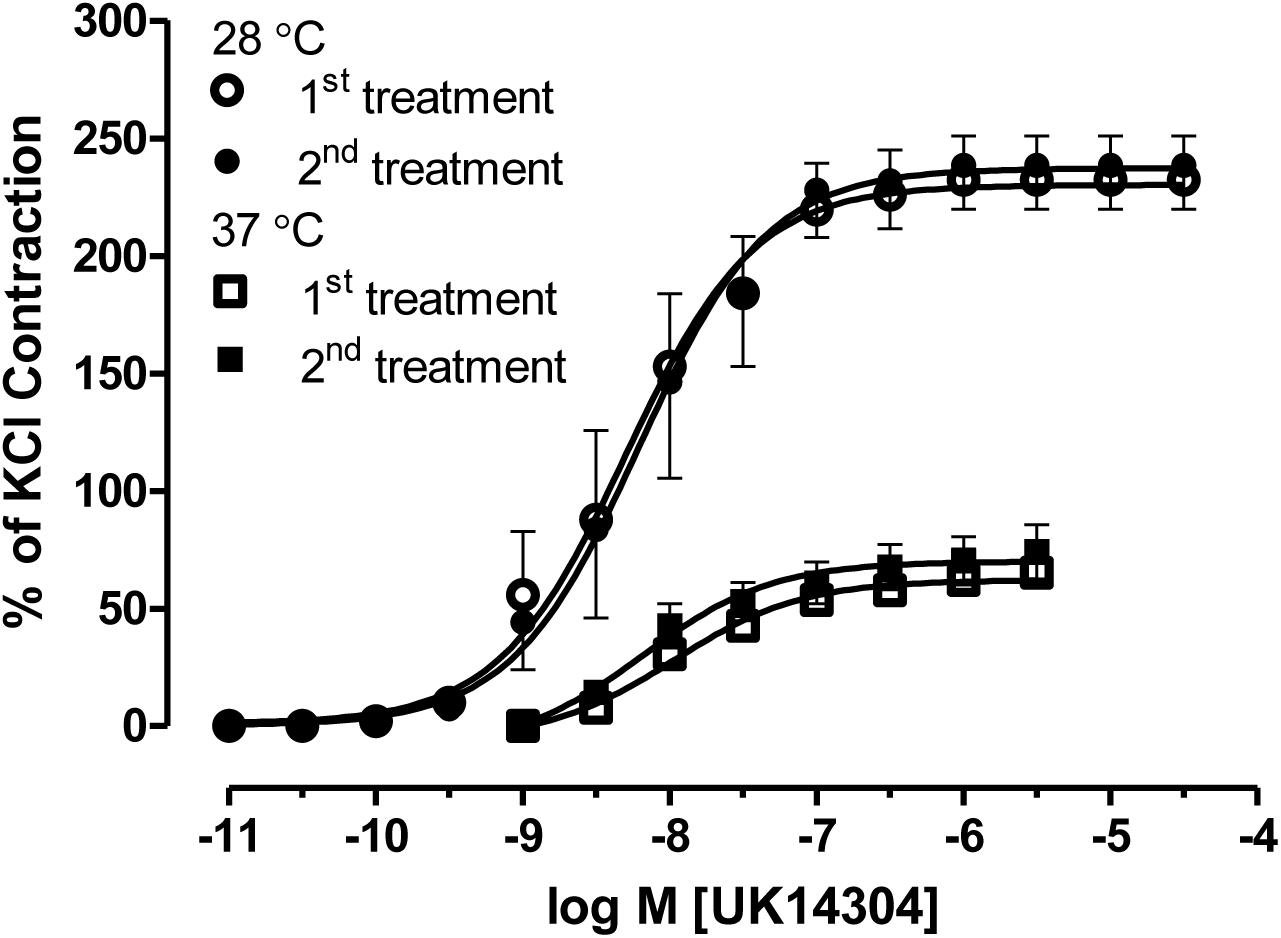
Effect of repeated treatments of UK14304 on CRCs of the rat tail veins. Contraction generated by the first administration of UK14304 is indicated as 1^st^ treatment and the second administration 1 hr later as 2^nd^ treatment at 37 °C and 28 °C are shown. Points are the mean ± S.E.M. of 4 experiments using individual rat tail veins, each taken from different animals.

### 3.7 Effect of moderate cooling on UK14304-mediated rat tail vein contraction

Rat tail serves as a thermoregulatory organ and anatomy of rat tail reveals that two lateral veins are cutaneous while mid-ventral tail artery lies in a groove which is closed by a ventral fascia (Flavahan, 1991). It is highly likely that vascular tone of later veins varies in accordance with temperature fluctuation. Effect of moderate warmup from 28 °C to 37°C on UK14304-mediated was studied first. CRCs for UK14304 generated at 37 °C and 28 °C were recorded; EC_50_’s as well as contraction produced at each concentration of UK14304 presented in tension (mg) (Fig. 6A) or normalized to percentage of KCl-induced contraction (Fig. 6B) were compared. Warm up to 37 °C rightward shifted CRC for UK14304 (*p*<0.001), with decreased mean EC_50_ and significantly inhibited vasoconstriction. Contractions measured in tension (mg) caused by most agonist concentrations, i.e., 1×10^−9^ M through to 3×10^−7^ M, were significantly greater at 28 °C (Fig. 6A). At high concentration of 1×10^−6^ M, vasoconstriction reached maximal capacity without further increase in tension at even higher concentration of UK14304. There was not a significant difference in contraction measured in tension (1043±142 mg at 28 °C vs. 690±88 mg at 37 °C, Fig. 6A). To compare contraction among different tissues, contraction measured in tension was standardized to 60 mM KCl-induced contraction. Normalized as percentage of KCl-induced contraction at 28 °C, warmup increased mean EC_50_ by 11-fold (1.6 ± 0.3 nM at 28 °C vs. 18 ± 7 nM at 37 °C, *p*<0.001) and significantly inhibited contractions at UK14304 concentration from 1×10^−9^ M through 1×10^−5^ M (Fig.6B; *, *p*<0.001). The maximal contraction normalized to KCl contraction at 28 °C was changed by 46 % (206 ± 12% at 28 °C vs. 141 ± 9 % at 37 °C). Experiments were also performed following a similar protocol but reversing the sequence of temperature change, i.e., cooling bath temperature from 37 °C to 28 °C. Cooling shifted CRCs for UK14304 leftward (*p*<0.0001, Fig. 6C), reduced mean EC_50_ by 8-fold (5±1 nM at 37 °C vs. 0.6±0.2 nM at 28 °C, *p*<0.001. Cooling significantly enhanced contractions at UK14304 concentration from 3×10^−10^ M through 1×10^−5^ M (Fig. 6C; *, *p*<0.001). The maximal contraction normalized to KCl contraction at 37 °C was increased by 47% (110 ± 6% at 37 °C vs. 162 ± 13% at 28 °C). As a normalization standard, contraction in response to 60 mM KCl was not altered by temperature shift (Fig. 6D). These experiments revealed cooling potentiated UK14304-mediated contraction of the cutaneous vein, irrelevant of order of temperature shift.

**Fig. 6.**
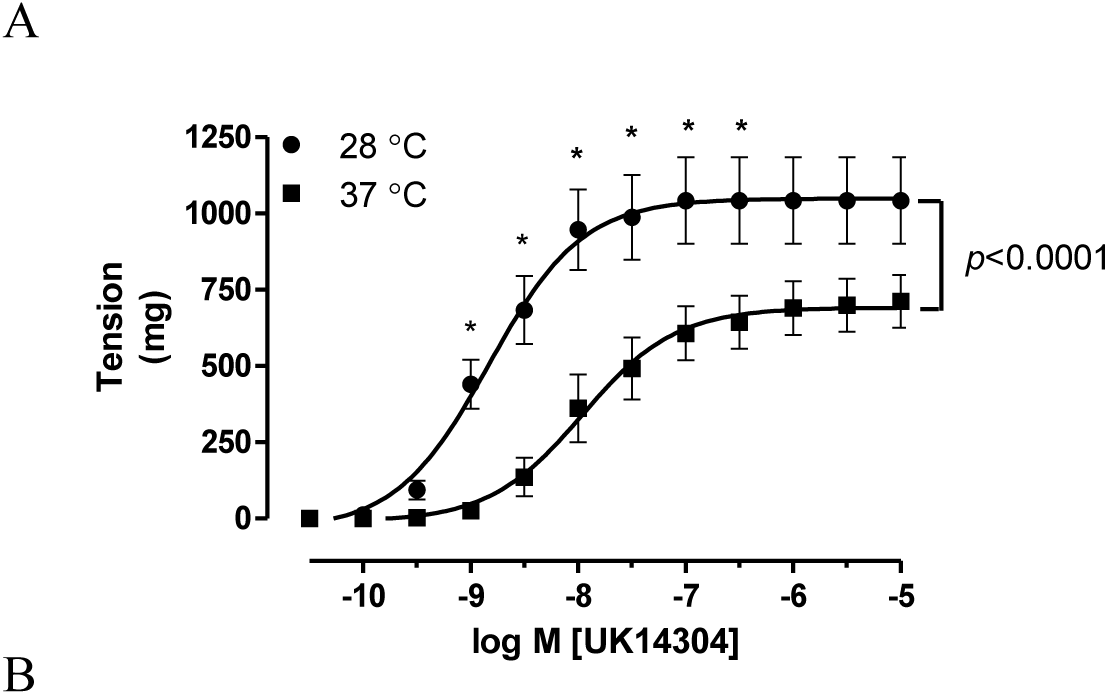
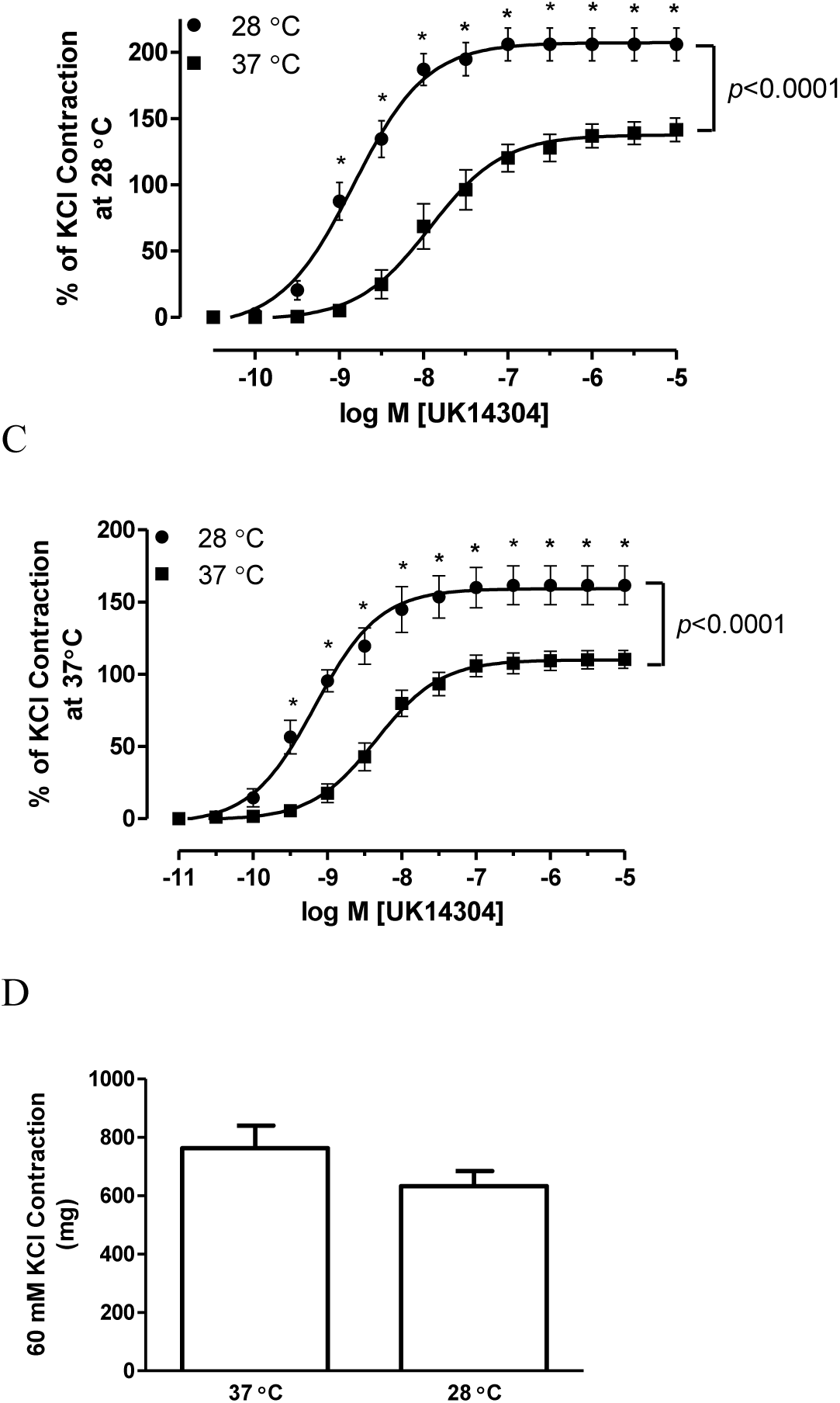
Effect of cooling on rat tail vein vasoconstriction in response to UK14304 and KCl. Mean concentration-response curves for UK14304-induced contraction of rat tail veins at 28 °C followed by increasing temperature to 37 °C is expressed as tension (A), or normalized as a percent of 60 mM KCl contraction produced at 28 °C (B). Contraction generated at 37 °C followed by decreasing temperature to 28 °C is expressed as a percent of 60 mM KCl contraction produced at 37 °C (C). Rat tail veins were contracted with 60 mM KCl at 37 °C and 28 °C and tension measured in mg (D). Points (A, B and C) and bars (D) are the mean ± S.E.M. generated from 6 individual experiments performed in duplicate or triplicate, with each experiment using veins from a different animal.

### 3.8 Effect of moderate cooling on another α_2_-agonist-mediated rat tail vein contraction

To determine whether the enhanced vasoconstriction after cooling was specific UK14304, CRCs for another α_2_-adrenoceptor agonist guanabenz were also generated at 28 °C and 37 °C. Guanabenz caused concentration-dependent contraction of the rat tail vein at both temperatures without necessity of precontraction by any other vasoactive agent. Similar to cooling effect on UK14304-mediated contraction of rat tail vein, CRCs for guanabenz at 28 °C were significantly different from those obtained at 37 °C (*p*<0.0001, Fig. 7) evidenced by significant decrease in mean EC_50_’s by 18-fold (2.5 ± 0.5 nM at 28 °C vs. 44 ± 17 nM at 37 °C, *p*<0.01) along with substantially increased contractile response to guanabenz in a concentration range from 1×10^−9^ M through to 1×10^−5^ M at 28 °C compared to 37 °C (*, *p*<0.01; Fig. 7). The maximal contraction was enhanced by 54% (96 ± 8 % at 37 °C vs. 147 ± 7 % at 28 °C). UK14304 Thus, moderate cooling to 28 °C enhanced cutaneous venous contraction are common to α_2_-adrenoceptor selective agonists.

**Fig. 7.**
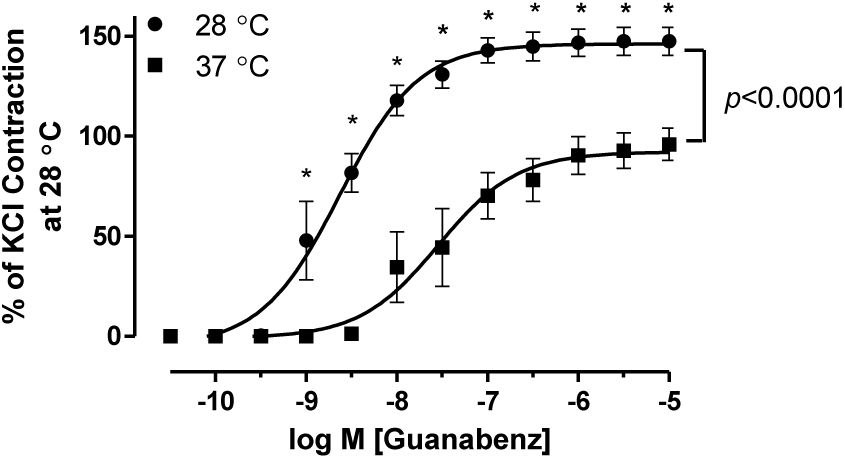
Mean concentration-response curves for guanabenz-induced contraction of rat tail veins. Contraction is expressed as a percent of KCl contraction produced at 28 °C. Points are the mean ± S.E.M. of 4 experiments using individual rat tail veins, each taken from different animals.

**Fig. 8.**
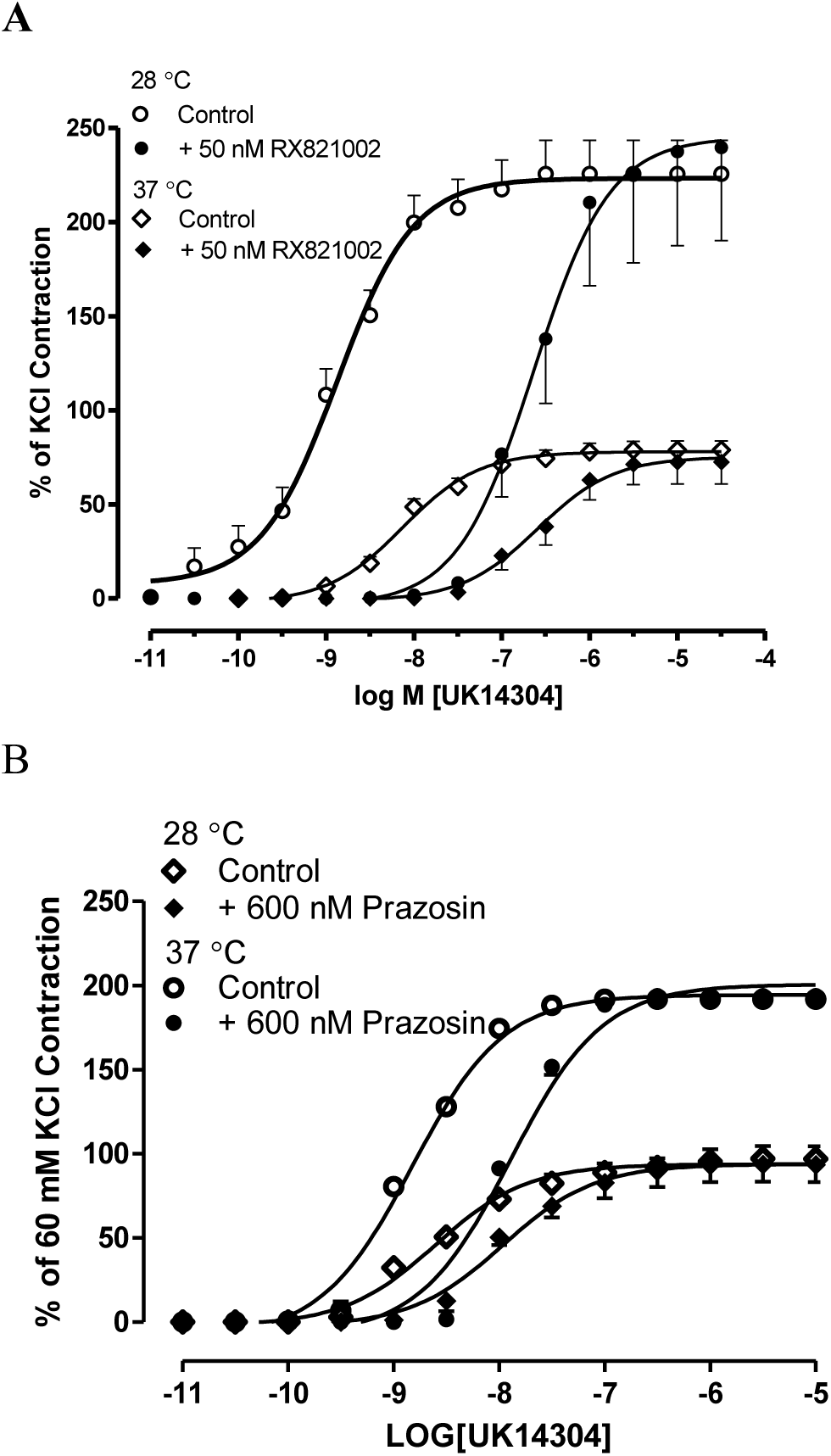
Antagonism against UK14304 concentration-response curves for contraction of rat tail veins by RX821002 (A) or prazosin (B) at 37 °C and 28 °C. Contraction to UK14304 is expressed as a percent of KCl contraction. Points are the mean ± S.E.M. of 3-4 experiments using individual rat tail veins, each taken from different animals.

### 3.9 Antagonism of α_2_-AR antagonist on UK14304–induced CRC

UK14304 is known to selective for α_2_-AR. To confirm UK14304 mediated venous contraction via α_2_-AR at both 37 °C and 28 °C, two antagonists with different selectivity were applied. RX821002 has been known to highly selective for α_2_-adrenoceptor for decades (Clarke & Harris, 2002; O’Rourke et al., 1994), while prazosin exhibited high selectivity for α_1_-ARs and much lower affinity for α_2_-ARs in rodents (Morrow & Creese, 1986; Uhlen, Xia, Chhajlani, Felder, & Wikberg, 1992). RX821002 and prazosin competitively antagonized UK14304-mediated venous contraction at 37 °C and 28 °C. Low concentration of potent α_2_-AR antagonist RX821002 (50 nM) reduced potency of UK14304 by 34- and 173-fold at 37 °C and 28 °C, respectively (Fig. 8A). Whereas high concentration of potent α_1_-AR antagonist prazosin (600 nM) rightward shifted CRCs by 5- and 8-fold at 37 °C and 28 °C, respectively (Fig. 8B), ruling out possibility of α_1_-AR involvement in UK14304-mediated contraction of rat tail vein.

## Discussion

Location allows unambiguous distinction of venous and arterial components of the circulation in this preparation. Use of arteries from rat tail has been reported (Jantschak et al., 2010; Souza, Padilha, Stefanon, & Vassallo, 2008). Venous tissue was easily accessible, however anatomy of tail lateral vein makes it obstacle to relocate lumen openning for mounting after blood drainage. The current report describes the reliable harvest of lateral veins without specialist resources, allowing prompt and easy retrieval of these vessels to be investigated by well established and reliable techniques such as organ bath. This study characterized vascular reactivity mediated through α_2_-ARs of the rat tail later vein. An underlying hypothesis of this study was that the tail later vein would response to α_2_-AR selective agnonists similarly to tail ventral artery or mouse tail artery. KCl-induced contraction is due to membrane depolarization causing the opening of voltage-gated Ca^2+^ channels, activation of Ca^2+^-dependent myosin light chain kinase and increase in myosin light chain phosphorylation, called electromechanical coupling (Ratz, Berg, Urban, & Miner, 2005). UK14304 or guanabenz induces venous contraction via activating α_2_-ARs, one of G-protein coupled receptors (GPCRs). GPCRs mediate signaling transduction through a variety of events including receptor activation, dissociation of trimeric G proteins and amplification of the signal through multiple cell messengers, called pharmacomechanical coupling. Due to its distinctive stimulus-response coupling mechanisms, KCl-induced contraction is independent of variations associated with GPCR activation. Tension induced by 60 mM KCl is commonly used as a standard to normalize the tension generated by GPCRs among different tissues. It was found that moderate cooling depressed KCl contraction of vascular smooth muscle in canine blood vessels (Rusch, Shepherd, & Vanhoutte, 1981; Tsukada & Chiba, 2000; P. M. Vanhoutte & Lorenz, 1970; P. M. Vanhoutte & Shepherd, 1970). Anatomy of tail lateral vein renders it high sensitivity to temperature fluctuation. Whether KCl contraction in rat tail lateral vein is susceptible to moderate cooling was examined. Tension produced by rat tail lateral veins in response to 60 mM KCl was not different at 37 °C compared to 28 °C.

The presence of endothelium in blood vessels may alter the contractile response to agonists. For example, removal of endothelium in mouse tail arteries impaired the constriction caused by activation of α_2_-adrenoceptors (Bailey, Mitra, Flavahan, Bergdall, & Flavahan, 2007). In rabbit ear arteries, contraction to noradrenaline, phenylephrine and BHT920 was endothelium-independent at 37 °C; whereas cooling to 24 °C increased contraction of rabbit ear arteries in the absence of endothelium (Garcia-Villalon et al., 1992). Presence of endothelium had not effect on contractile responses to UK14304 in rat tail lateral vein at 37 °C and 28 °C.

Desensitization may occur to α_2_-AR following prolonged or multiple treatments with agonist. This study showed that CRCs for UK14304 remained same after repeated administration of UK14304 at 37 °C and 28 °C, ruling out likelihood of desensitization. As such, each venous ring was safe to serve as internal control in studies.

Distal segments of mouse tail artery exhibited greater vasoconstriction in response to α_2_-adrenoceptor activation compared to proximal segments (Chotani et al., 2000). Therefore, the effect of the distance from the torso of the rat tail vein on the contractile response to UK14304 was also examined. At 37 °C, venous sections close to or away from torso exhibited uniform contractions whereas contractile profile generated in the distal portion was significantly different from that in the proximal portion at lower temperature of 28 °C with no significant changes in either potency or maximal contraction.

In contrast to weak contraction in response to α_2_-ARs agonists without appropriate pre-contraction or insensitive to decreasing temperature in the rat tail ventral artery (Harker, Ousley, Bowman, & Porter, 1991; Jantschak et al., 2010; Savino & Varela, 1999), the later vein robustly responded promptly to α_2_-AR selective agonists UK14304 and guanabenz at both tested temperatures 37 °C and 28 °C. Rat tail lateral vein exhibited enhanced vasoconstriction with higher potency activated by α_2_-adrenoceptor selective agonist UK14304 or guanabenz at moderate cooling temperature of 28 °C, independent of temperature shift order. Two α_2_-AR selective agonists tested in the study revealed the potentiated venous vasoconstriction observed at 28 °C was not unique for UK14304. It should be noted that this effect was also showed in other α_2_-adrenoceptor selective agonists BHT933 and dexmedetomidine (data not showed). Antagonism against UK14304 CRCs at 37 °C and 28 °C by high concentration of α_1_-AR selective antagonist prazosin and low concentration of α_2_-AR selective antagonist RX821002 validates the principal role of α_2_-AR. All these findings collectively suggest α_2_-AR rather than any agonist leads to significantly potentiated vasoconstriction at moderate cooling temperature.

Both mouse tail arteries and rat tail arteries have been well characterized, demonstrating differentiated response to temperature fluctuation. Nevertheless, studies on tail veins were not yet reported. Isolation of mouse tail veins as an *ex vivo* model is still beyond currently available techniques, although injection into blood drawn from mouse tail vein have become common in research. Rat tail vein is much easily accessible in dissection compared to its arterial compartment, rendering it a potential candidate as an *ex vivo* model worthy of being characterized. Vascular tone mediated through α_2_-AR in rat tail vein was first examined in our laboratory. The rat tail vein can serve as an appropriate *ex vivo* model to study vascular reactivity of cutaneous vessels under physiologic and/or pathophysiologic states such as Raynaud’s phenomenon.

## Abbreviations

AR: Adrenergic receptor
CRC: concentration-response cur

